# Unequal access to resources undermines global sustainability

**DOI:** 10.1101/2020.07.27.222414

**Authors:** Kirsten Henderson, Michel Loreau

## Abstract

We are a global society with various land use patterns, inequality, and the movement of people and goods. The various practices and behaviours associated with our current society raise questions about the future sustainability of the human population and the natural environment. We derive a model of a global socio-ecological system to explore the connections between human well-being and land resources, specifically looking at resource accessibility, conservation initiatives and human migration between two economically diverse regions. We find that the spatial aspect of a global system with two distinct regions, allows for faster development of technology, higher peaks in population size, greater natural land degradation, and generally speaking lower population well-being in the long-term. The unequal access to resources and differences in technological progress, alter the outcome of land management (i.e., conservation) and social behaviours (i.e., migration). We conclude that any socio-ecological management practices should be conscientious of the diversity in land access, population size, population well-being and development within the global society, as the potential for unintended consequences is high. Inequality needs to be addressed to promote sustainability.

## 1 INTRODUCTION

Goodhart’s Law states that “when a measure becomes a target, it ceases to be a good measure” (Chrystal et al., 2003). This statement is particularly relevant to our global socio-ecological and economic system, where the pursuit of well-being and economic growth can cause people to overlook other aspects of life, such as biodiversity and ecosystem services. Therefore, efforts to promote environmental sustainability and population ‘well-being’ need to consider the entirety of the socio-ecological system, as ecosystem services are essential to wealth, well-being, and sustainability (Costanza et al., 2014).

More often though the environment is valued only for the productive assets (i.e. resources), which leads to inequality and poor land management. Inequality is driven by different land-use practices and investment choices that fail to distribute resources equally (Coomes et al., 2016). Inadequate distribution forces more land to be converted, which can lead to a cycle of poor land management and social inequality and pushes development away from environmental sustainability (Hasegawa et al., 2019; Boyce, 1994; Cumming and von Cramon-Taubadel, 2018). Furthermore, there exists a positive feedback between power and wealth, which reinforces inequality, such that in a finite system when one benefits and the other loses, the result of applying random processes is extreme inequality (Scheffer et al., 2017).

Modern practices are built on the idea that wealth and development of knowledge can continue infinitely (Cass and Mitra, 1991), which requires that the pace of population growth increases with social organization, such that development is not allowed to stagnate (Bettencourt et al., 2007). If technological growth does not continue, economic expansion will increase the demand on the ecological system (Clow, 1998). However, Cumming and von Cramon-Taubadel (2018) found that the development of countries is not sufficient for environmental sustainability, as under the current two-economy system (i.e., high- and low-income) both economies are not allowed to continue developing or cannot simultaneously accumulate wealth (Cumming and von Cramon-Taubadel, 2018) – the less developed economy inevitably experiences poor living conditions.

When living conditions become undesirable, it becomes beneficial for individuals to move. Sweden in the 19th century experienced mass movements of people, which has been attributed to poor resource availability and accessibility (Clarke and Low, 1992). In the North of Sweden, where the land was less productive and the carrying capacity was minimal, experienced the highest number of individuals leaving the region. Likewise, drought has been a major factor in temporary and indefinite migration, by altering settlements and agricultural practices, as observed during the Dust Bowl of the 1930s in North America, and the severe droughts in Africa through 1980s and 1990s (McLeman, 2014). Indeed, migration has been shown to allow individuals to inhabit less favourable environments through temporary dispersal (Holt, 2008) and even has the ability to reduce poverty by moving to regions with more opportunities or wealth (Adams Jr and Page, 2005). Observational evidence shows that migration is the result of many factors, but the basic theory is that either the local conditions are insufficient, forcing people to leave, or the conditions elsewhere are comparatively better than the local conditions, attracting new individuals (Grigg, 1977).

Among the many social factors that influence dispersal — policy, family, job opportunities (Gonzalez et al., 2008) — income inequality can have the largest impact, both directly and indirectly. As mentioned above inequality leads to greater land degradation, and severe land degradation forces people to disperse. This phenomenon is more likely to affect low-income individuals, for which agriculture is the main income generator (Levy and Patz, 2015). However the paradox of migration is that the cost is too high for the poor to disperse (Black et al., 2011) and the wealthy do not benefit from dispersing (Towner, 1999). If people are unable to move and the land is degraded, they will inevitably be embroiled in poverty and experience poor well-being (Barbier and Hochard, 2016).

Here we build and analyse a model to explore land and social dynamics in space. We generate a model consisting of two regions with inequality, through differences in access to technology and resources. We compare the ‘real-life’ model to a uniform one-region system, in addition to scenarios that alter income status within a region and dispersal between regions. Furthermore, we incorporate conservation and restoration practices in the two-region system with distinct populations and practices, hypothesizing that increasing the natural area can contribute to a sustainable and desirable future for humanity.

## 2 BRIEF MODEL DESCRIPTION

Previous work has modelled the relationship between differing economies (e.g., Human Development Index 1 (HDI1) regions and HDI4 regions) and distinct practices (Cumming and von Cramon-Taubadel, 2018). This is supported by empirical data showing that there are two groups of individuals with distinct demography, development structures, and consumption patterns (Oswald et al., 2020), in addition to an analysis of The World Bank (2019a) data given in the appendix. We use this idea of distinct economies with distinct practices and apply it to an ODE model of global land management and population growth (Henderson and Loreau, 2019). We modified the Henderson and Loreau model to incorporate two regions, movement of people and goods and inequality. The model simulates a simplified global system with two regions and two subpopulations within each region. The regions represent higher income and lower income economies and development structures (*j* = *L, H*), each with subpopulations that are also classified as higher income and lower income (*P*_*i, j*_, where *i* = *L, H* represents the population income level, and *j* = *L, H* reflects the region income level). The higher income region refers to a GDP above the global average and the lower income region refers to a GDP below the global average, we have included a spreadsheet with this data in the appendix. The subpopulations LH and HL reflect the middle income groups in the ‘real world’. These distinctions were made on the basis that there are clear differences in the stages of the demographic transition, income, social norms, land-use practices, consumption habits and technological development between regions and populations classified as HI or LI. Kernel density plots are given in the appendix to show distinct groupings for higher and lower income regions, with more ambiguous differences between middle income groups. The four subpopulations in the model represent the spectrum of income groups globally, showing the variation in consumption levels, birth rates, death rates, research and development expenditure, and resource production. Equations and a full description of the model are provided in the appendix.

### 2.1 Human population

The population growth function, which takes into consideration recruitment and adult mortality rates, is dependent on resource accessibility (ha/pers., which is calculated as a function of technology and land capacity). When population growth is plotted against resource accessibility we see a non-monotonic curve that increases initially with resources and then declines as resource accessibility surpasses the basic needs level. The details of this theory are described in Henderson and Loreau (2019). Resource accessibility also moderates the rate at which individuals change income status. Once an accessible resource threshold (ha/ind.) is crossed — determined by World Bank income classifications (The World Bank, 2019a) and the ecological footprint of each country (Global Footprint Network, 2019) — individuals can become higher income or lower income. The shift in status increases exponentially with resources, when individuals shift from lower to higher income; and the shift in status decreases logistically from higher to lower income. Furthermore, individuals are able to move from one region to another by comparing the accessible resources in the foreign region with their own resource accessibility. In the model, a sigmoidal curve is used to represent the relationship between resource accessibility and dispersal.

### 2.2 Land cover

The two regions are composed of natural land (*N*_*j*_), where natural land describes ‘semi-natural’ and natural land, such as grasslands, tree-covered areas, shrub-covered areas (the full list of natural land areas, as described by the FAO, is provided in the appendix); agricultural land (*A*_*j*_), which is described as croplands by the FAO; and unused land (*U*_*j*_), which is the total land area (*L*_*j*_) minus *A*_*j*_ and *N*_*j*_. Unused land describes all land that is not agricultural or natural, such as urban, degraded land, and minimally productive land (i.e., glaciers, barren land). Landuse practices include local and foreign use of land from degradation and cross-degradation to agricultural conversion, restoration (human and natural regeneration) and the option to include conservation.

The function for degradation and consumption of *N*_*j*_ and *A*_*j*_ (*j* = *L, H*) describes a linear dependence on the population size, the demand for resources and the share of land used by the local population. The remaining proportion of land not used by the local (*j*) individuals may be consumed and degraded by the individuals from the foreign region (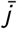, where 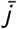 is the opposite of *j*, such that if *j* = *L*, 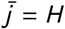 and vice versa).

Conversion from *N*_*j*_ to *A*_*j*_ depends on the demand from the population (*P*_*j*_) and technology (*T*_*j*_), in each region. When technology is abundant agricultural yield increases without the need for more land conversion. Furthermore, it has also been shown that agricultural production improves with surrounding natural land area (Bennett et al., 2009; Braat and De Groot, 2012). Therefore, agriculture degradation is a function of consumption and natural degradation, which are moderated by the supply of ecosystem services. Abundant natural land cover (*N*_*j*_, *C*_*j*_) buffers agricultural land degradation, which is reflected by an exponentially decreasing function in the model (details in the appendix).

Restoration is a function of both the natural and human processes of converting unused or degraded land (*U*_*j*_) back into natural land (*N*_*j*_). Conservation in the model refers to a fraction of natural land set aside, which provides individuals and the local environment with non-provisioning ecosystem services. Conservation occurs at a constant rate that is bound by the proportion of desired conserved land and already existing natural land.

### 2.3 Technology & development

Technology and development are major drivers of population dynamics via resource accessibility. It is estimated that higher income regions are more developed than lower income regions, in terms of education, medicine, machinery, etc. (The World Bank, 2019a; Kummu et al., 2018; Sarkodie and Adams, 2020; Sen and Laha, 2018). Therefore, we include two technology variables, one for each region (*T*_*j*_, *j* = *L, H*) with different growth rates. The technology growth curve is a function of population size and density, and the availability of natural resources (i.e., *N*_*j*_ and *C*_*j*_). Technology has been shown to increase with population density, however there becomes a point where the number of individuals exceeds the capacity of natural land and limits the future development of technology (Clow, 1998).

Technology is a major determinant of power and resource accessibility – determining who will use what land, when, and how. We assume that technology builds upon itself, therefore the region with more technology has the potential to develop new technologies more quickly, akin to the power cycle described by Scheffer et al. (2017).

### 2.4 Resource acquisition

Resource accessibility per individual is dependent on the power they exert (a combination of technological development and population size, details in the appendix), the availability of agricultural and natural resources, the ability to acquire such resources, and the potential to enhance production yield with technology.

### 2.5 Model analysis and simulations

We first build a business as usual (BAU) model that uses historical trends over the last 260 years to simulate current population and land dynamic, we validate our findings with data from the World Bank Group given in the appendix. We then apply alternative land management practices (i.e., conservation in the LI region, conservation in the HI region, restoration) and social policies (i.e., migration, income status) to current trends and simulate the results over the next 740 years, at which point the results reach a sustained value (an analytic equilibrium is not possible). The model contains 12 variables, which makes finding an analytic solution for equilibrium values difficult. Furthermore, when discussing population dynamics, the shortterm, transient dynamics are generally of most interest (Ezard et al., 2010). We run the model long-term to give an idea of possible trends, but this does not necessarily infer an equilibrium or give quantitatively realistic results. The ODE model was run through MATLAB using odesolver 113. Parameter values, initial conditions and a range of scenario parameters are given in the appendix.

The restoration scenario involves the active conversion of degraded or unused land (*U*_*j*_) back into natural land (*N*_*j*_) by individuals in the local region (*P*_*j*_, includes both subpopulations within the region). There is a minimal natural conversion and a minimal anthropogenic conversion, however this scenario also looks at restoring natural land at 50 to 100 times the natural rate of restoration. By contrast, conservation is used to describe natural land (*N*_*j*_) being set aside — taking *N*_*j*_ and maintaining it in the conserved state (*C*_*j*_), such that individuals and land cover are provided with non-provisioning services, but the land is not available for harvest or manipulation. We vary the rate of conservation in a effort to find a link between sustainability and conserved land. Conservation is applied to the LI region alone, the HI region alone, and both regions together.

The no status change scenario looks at the impact of keeping individuals in their respective subpopulation, regardless of the acquired resources to which they have access. We also increased the rate of change between income groups, allowing individuals within each subpopulation to transition more quickly between income groups. Finally, for the migration scenarios, we prevented individuals from relocating to a different region. Additionally, we doubled the rate of migration to see how allowing more people into foreign regions would impact the socioecological system.

In addition, we compare the two-region system, with four subpopulations to a one-region system, with two subpopulations, to understand the role of the spatial distribution of land and people in the dynamics of our global system.

The individuals in the population are assigned a well-being based on the number of accessible hectares per person: famine is defined as having less than 0.55 ha/pers; poor well-being occurs when there are between 0.55 and 1 ha/pers.; moderate well-being is defined by accessible resources between 1 and 2 ha/pers.; good well-being is defined as accessible resources between 2 and 5 ha/pers.; excessive well-being is equivalent to more than 5 ha/pers.

## 3 RESULTS & DISCUSSION

### 3.1 Business as usual scenario

The model is able to regenerate human population and land cover patterns observed over the last 260 years, (*N*_*L,H*_ ≈ 0.5 * *L*_*L,H*_, *A*_*L*_ = 0.84 Bha, *A*_*H*_ = 0.64 Bha) and population size in each region (*j*) (*P*_*L*_ = 5.9B, *P*_*H*_ = 1.4B), using parameters estimated from historic data and theories on technology, demography and ecology. Furthermore, the model projections fit within the range of the predicted population numbers in 2070 (9.9 to 11.2B) from the United Nations Population Division (2019). Our higher income population in 2070 is 4.3B, which is on the upper end of the UN range for high- and upper-middle-income populations (2.75 to 4.2B); and our lower income population is estimated to be 7.3B, on the high end of the UN range for low- and lower-middle-income populations (5.7 to 7.2B). Generally, the population is still growing slowly in the year 2100. Unlike, the UN projections, the model described here runs after 2100, after which the model shows major changes in population dynamics driven by the spatial distribution of people and goods.

The model predicts three stages of population dynamics, based on resource accessibility (i.e., land cover and technology) and dispersal trends. The first 350 years (approximately from 1760 until 2100) are governed by resource accessibility which allows the population to grow, prior to natural land limitations; hence, why many scenarios are similar over this time span.

However, afterwards the access to resources changes the spatial distribution of individuals, and at this stage dispersal becomes the main driver. The subpopulations are reconfigured into different income groups, regions and population numbers as a result of the feedbacks between resource accessibility and dispersal, which causes a second wave of population growth. This alters technological development and degradation patterns, which ultimately impacts population growth and well-being. After 2100, dispersal becomes the primary driver of human population dynamics, and thereby also land cover patterns.

Finally, in 2250, the population starts to decline, as technology has long since stagnated and resource availability declines below adequate levels to maintain the human population. In the long-term, the BAU scenario leads to famine in the LI region and poor well-being in the HI region. Both regions experience a population decline, as a result of high death rates and little or no recruitment.

### 3.2 Impact of technology

The major differences between the two regions (higher income and lower income) can be attributed to the population size and technological development in each region. In general, technology allows the population to sustain a high well-being lifestyle, which contributes to a declining population and leads to minimal population numbers. Model simulations suggest that technology plateaus in 2070, as a result of declining natural land, which limits resource accessibility – forcing either well-being or human population size to decline, in the short-term. In the long-term, both decline. Without resources, technological development becomes difficult.Once the population and resources are sufficiently low or there is a mismatch between population density and resource availability, technology also begins to drop.

The higher income region has a technological advantage over the lower income region that ensures the higher income region has higher well-being and more access to resources than the lower income region. However, lower income populations produce people power and without the flow of people from the LI region to the HI region, technological development curtails in the higher income region. The model suggests that it is difficult for the lower income region to match the technological development of the higher income region, when resources from the LI region are being used by the HI region and the higher income subpopulation of the LI region seek opportunities in the higher income region. This resonates with Richardson (1995) stating that globalization of the economy has contributed to a rise in inequality.

### 3.3 Impact of dispersal

In addition to technology, dispersal is another key driver of the system. The model clearly shows that dispersal of individuals alters technological development, degradation patterns, and growth patterns. As mentioned above the second stage of the population trends observed in this model is governed by dispersal. Hence without dispersal from one region to another (Fig. 2), the population in the higher income region shrinks (*P*_*H*_ < 0.1B in 2750). There is not enough replacement growth within the HI region and without input from the lower income region, the population is small and declining. The direct effect of not allowing individuals to move from one region to the other, results in declining populations: one from excessive well-being and no population regeneration (HI region); the other from poor living conditions and high mortality rates. Doubling the dispersal rate has no qualitative differences to the BAU scenario.

**FIGURE 1.**
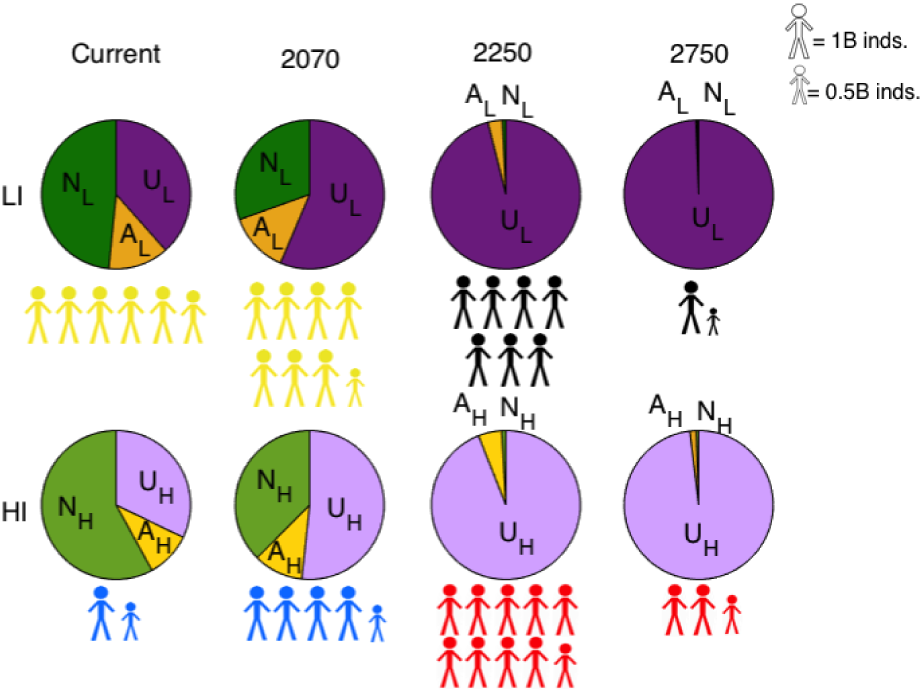
Business as usual scenario – land and population patterns. Currently natural land occupies the greatest area in both regions (*N*_*L*_, *N*_*H*_), followed by unused land (*U*_*L*_, *U*_*H*_) and approximately 10% agricultural land (*A*_*L*_, *A*_*H*_). By 2750, *U*_*L*_ and *U*_*H*_ remain, with negligible fractions of *N*_*L*_, *N*_*H*_, *A*_*L*_ and *A*_*H*_. The population (*P*_*j*_) in each region, *j* (*j* = *L, H*), is represented by the number of stick figures and shrunken stick figures represent fractional billions. The population peaks after 2100, while the well-being peaks in 2070. The well-being (*W*_*L*_, *W*_*H*_) is determined by the accessible resources per person (ha/pers.): yellow = moderate well-being, blue = excessive, black = famine, red = poor.

**FIGURE 2.**
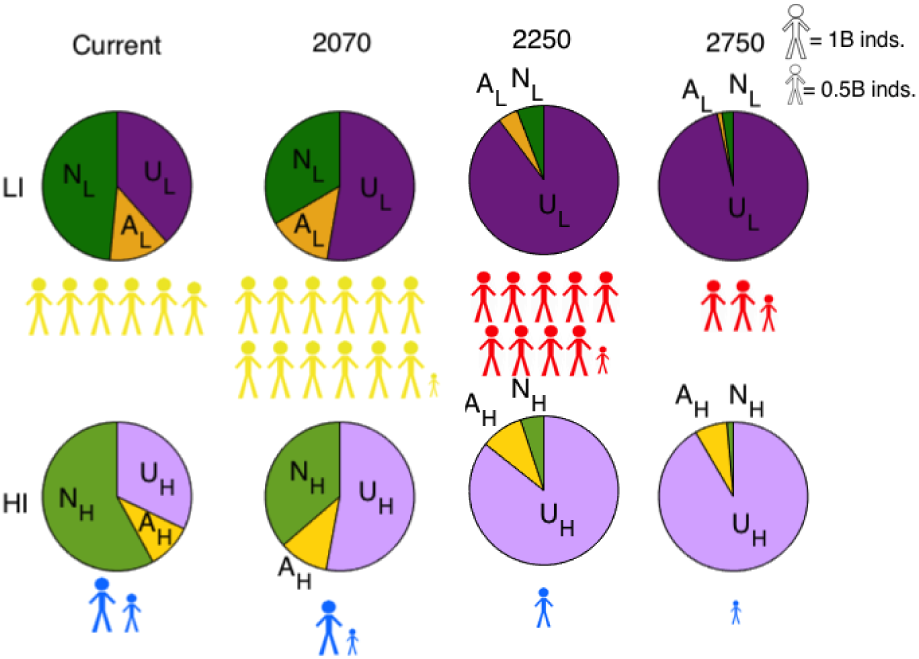
No dispersal scenario – land and population patterns. The population in the HI region (*P*_*H*_) is low compared to BAU population (current-2750), where *P*_*H*_ < 0.1B inds. exist with excessive well-being (blue), 0.46 Bha of agricultural land (*A*_*H*_) and minimal natural land (*N*_*H*_ < 0.1 Bha) in 2750. The population in the LI region more than doubles between now and 2070 (*P*_*L*_ = 12.1B inds.), maintaining a moderate well-being (yellow). However, as resource accessibility diminishes, so does the population size (from a lack of resources) and well-being. By 2750, there are 2.5B inds. with a poor well-being and a sliver of natural land remains until 2750 (*N*_*L*_ = 0.16 Bha).

Based on the model feedbacks and UN reports (UNnews, 2019), dispersal is a reaction to insufficient resource accessibility, whether relative or real. From model simulations we infer that individuals seek better opportunities, which results in short-term increases in resource accessibility and growth. Yet, in the long-term leads to homogeneously poor well-being, if there is no change in consumption habits. We deduce that dispersal temporarily masks or dilutes feedbacks between resource accessibility and population dynamics. As a result, dispersal encourages populations to grow beyond the resource accessibility at the regional level, by allowing individuals to move and access more resources elsewhere.

### 3.4 Conservation scenario

Conservation in a region (*j*), results in massive dispersal out of the other region (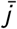) into the region (*j*), with conserved natural land (natural land that does not provide any provisioning services to the human population, *C*_*j*_, Fig. 3). The region with conservation experiences increases in growth and dispersal, as individuals from the non-conservation region flow in, causing changes in resource accessibility. Initially, the fluctuations in resource accessibility promote growth; however, as the population grows the resource accessibility per capita declines significantly and causes a decline in population.

**FIGURE 3.**
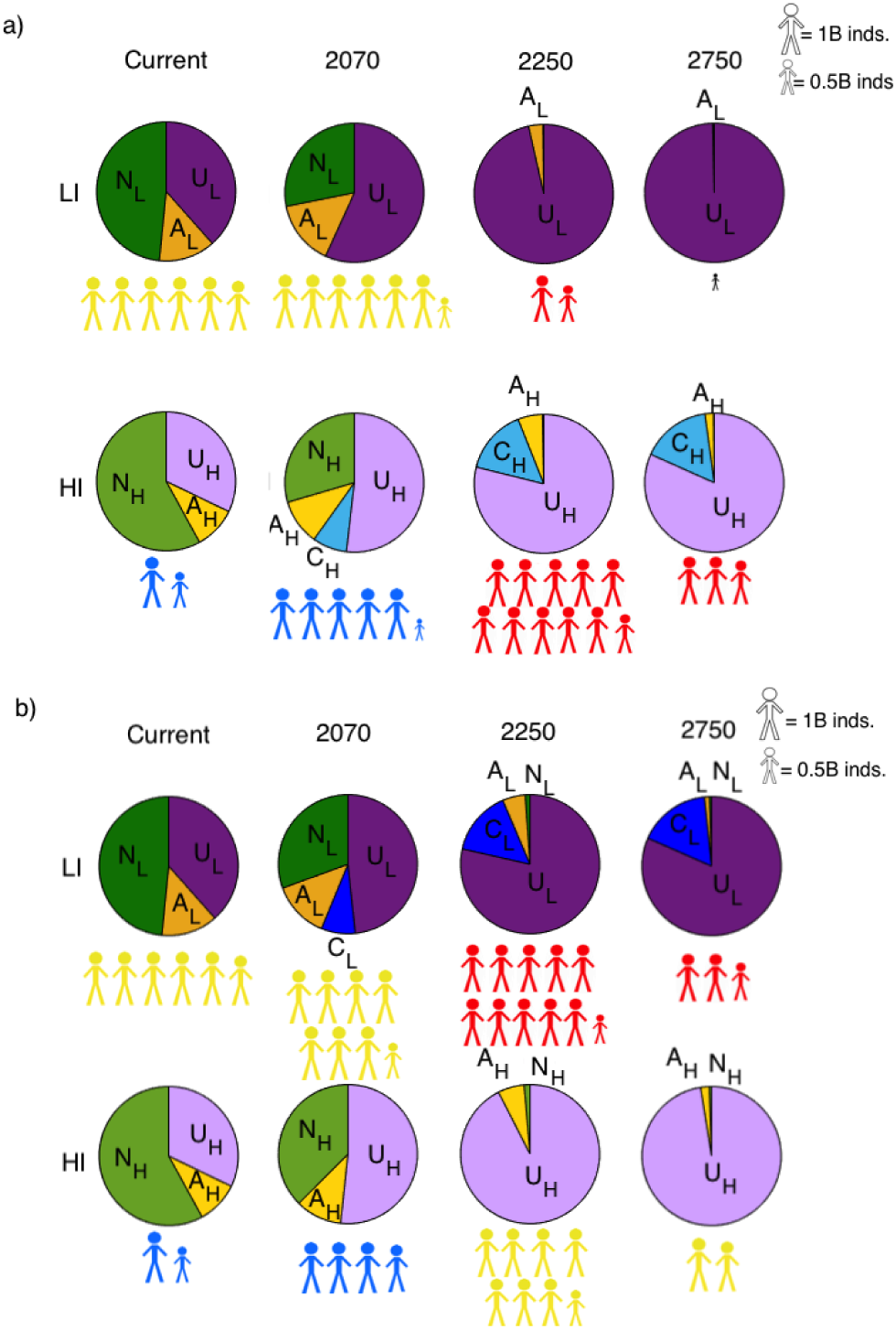
a) HI Conservation – land and population patterns. Conservation entails setting aside a proportion of natural land for ecosystem services, excluding provisioning services (*C*_*H*_). *C*_*H*_ eclipses natural land, as *N*_*H*_ becomes *C*_*H*_. *C*_*H*_ maintains more agricultural land in 2750 (*A*_*H*_ = 0.13 Bha), but has no impact on the land dynamics of the LI region (*N*_*L*_, *A*_*L*_). Conservation in the HI region, causes a decline in the population in the LI region (*P*_*L*_ = 0.1 in 2750) and a famine state (*W*_*L*_, black). Initially, the population in the HI region increases (*P*_*H*_ = 10.7B inds. in 2250) and then decreases due to a lack of resources (*W*_*H*_ = *poor*) over a prolonged period (2250 to 2750). b) LI Conservation – land and population dynamics. Conservation in the lower income region reduces the long-term unused land fraction (*U*_*L*_) by replacing it with conserved land (*C*_*L*_). The other land uses – natural land (*N*_*L*_, *N*_*H*_) and agricultural land (*A*_*L*_, *A*_*H*_) – are minimal, yet still higher than the BAU scenario. Both populations in the HI and LI regions have a higher well-being compared to the BAU scenario in 2750 (*W*_*H*_ = *moderate, W*_*L*_ = *poor*). The population size is similar to the BAU, however the number in each region is flipped; *P*_*H*_ = 7.9B and *P*_*L*_ = 10.3B in 2250, *P*_*H*_ = 1.8B and *P*_*L*_ = 2.5B in 2750.

In the higher income region, when conservation is applied (Fig. 3a), the natural land cover (natural land that is available to individuals for provisioning services, *N*_*j*_) is similar to the BAU scenario, however the amount of degraded or unused land declines (*U*_*j*_) by at least 1 Bha.

When conservation is applied to the lower income region (Fig. 3b), there remains a minimal quantity of natural land (*N*_*L*_, *N*_*H*_) and agricultural land (*A*_*L*_, *A*_*H*_), in both regions. When more individuals move to the LI region, as is the case when conservation is introduced, the wellbeing increases in both regions (*W*_*L*_, *W*_*H*_), due to lower consumption rates in the LI region. The sustained technology value in the LI region is greater (*T*_*L*_ = 3.3).

The one-region conservation scenario provides an interesting contrast to the two-region system. Conservation in the one-region system generates the greatest abundance of natural land, in the conserved state, of all scenarios. Whereas, in the two-region system, agricultural land increases and there is an increase in natural land, in the conserved state. In the higher income one-region system conservation has no direct or indirect outcome on the human population, the land is merely shifted from unused to conserved nature. This is similar to the case of no migration or conservation in all regions in the two-region case. Empirically, the regions where dispersal is unlikely, for social or economic reasons, with populations that are highly dependent on the local environment, conservation policies can be restrictive. This is consistent with the Cazalis et al. (2018) model, where extreme conservation lead to famine, and our oneregion, lower income, conservation scenario (details in appendix). For example, small-scale subsistence farmers, with minimal income or technology may experience detrimental consequences from strict conservation policies. This has been observed in Nepal (Brown, 1998), however conservation designed to help subsistence farmers has benefited yields in Ethiopia (Bekele, 2005).

### 3.5 Restoration scenario

Restoration increases natural land (*N*_*L*_, *N*_*H*_) and agricultural land (*A*_*L*_, *A*_*H*_) area in both regions (Fig. 4). Unlike most scenarios, restoration maintains *N* until 2750 (*N*_*L*_ = 1.34 Bha, *N*_*H*_ = 1.74 Bha). With an increase in land cover there is an increase in resource accessibility, which allows the population (*P*_*L*_, *P*_*H*_) to continue growing along with technology (*T*_*L*_, *T*_*H*_). However, this relatively unchecked population growth in both regions leads to very high populations (*P*_*L*_ = 43.4B, *P*_*H*_ = 67.3B) with a poor well-being, in the long-term. Technology reaches a maximum of *T*_*L*_ = 6.9 and *T*_*H*_ = 24.5, the highest of all scenarios. The land is continuously converted back to natural land, which prolongs the amount of time before technology is limited by the imposed natural land threshold (*N*_*th*_).

**FIGURE 4.**
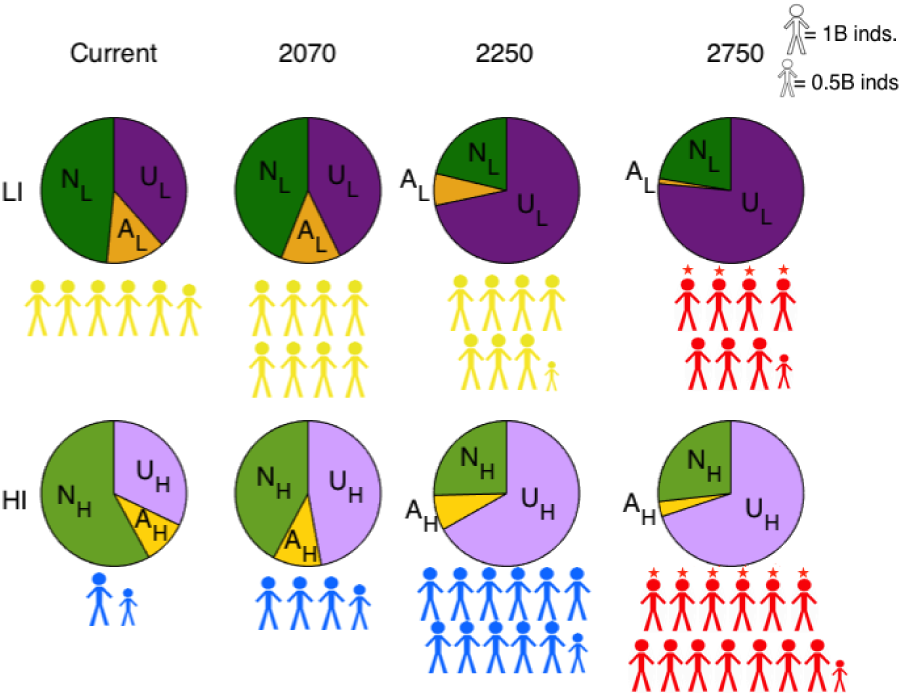
Restoration – land and population patterns. Restoration increases the sustained area of natural land (*N*_*L*_ ≈ 1.3, *N*_*H*_ ≈ 1.7), which drives an increase in population size, in both regions. The population well-being is maintained at its current state until 2250 (*W*_*L*_ = moderate, *W*_*H*_ = excessive), after which the population becomes too large (*P*_*H*_ = 67.3B inds., *P*_*L*_ = 43.3B inds.) for the availability of resources (*N* and *A*).

Restoration diminishes the rate of dispersal for the first 600 years (until 2350) in the higher income region, as the HI region benefits from the influx of resources and has no reason to disperse. However, we infer from the simulations that the lower income region population uses these newfound resources to seek better opportunities. As a result, the population increases in both regions, all populations disperse, looking for more resources, only to find that resources are limited globally.

Dispersal also subverts attempts to restore natural land and increase well-being. Restoration in the one-region case (see below) benefits the population well-being, without increasing the population size (Fig. 5c). The same is not true of the two-region, ‘real-world’, case. If population growth and well-being stagnated in 2250, restoration would be beneficial to natural land recovery and population dynamics. However, without a change in habits, restoration encourages rapid growth and poor well-being, long-term.

**FIGURE 5.**
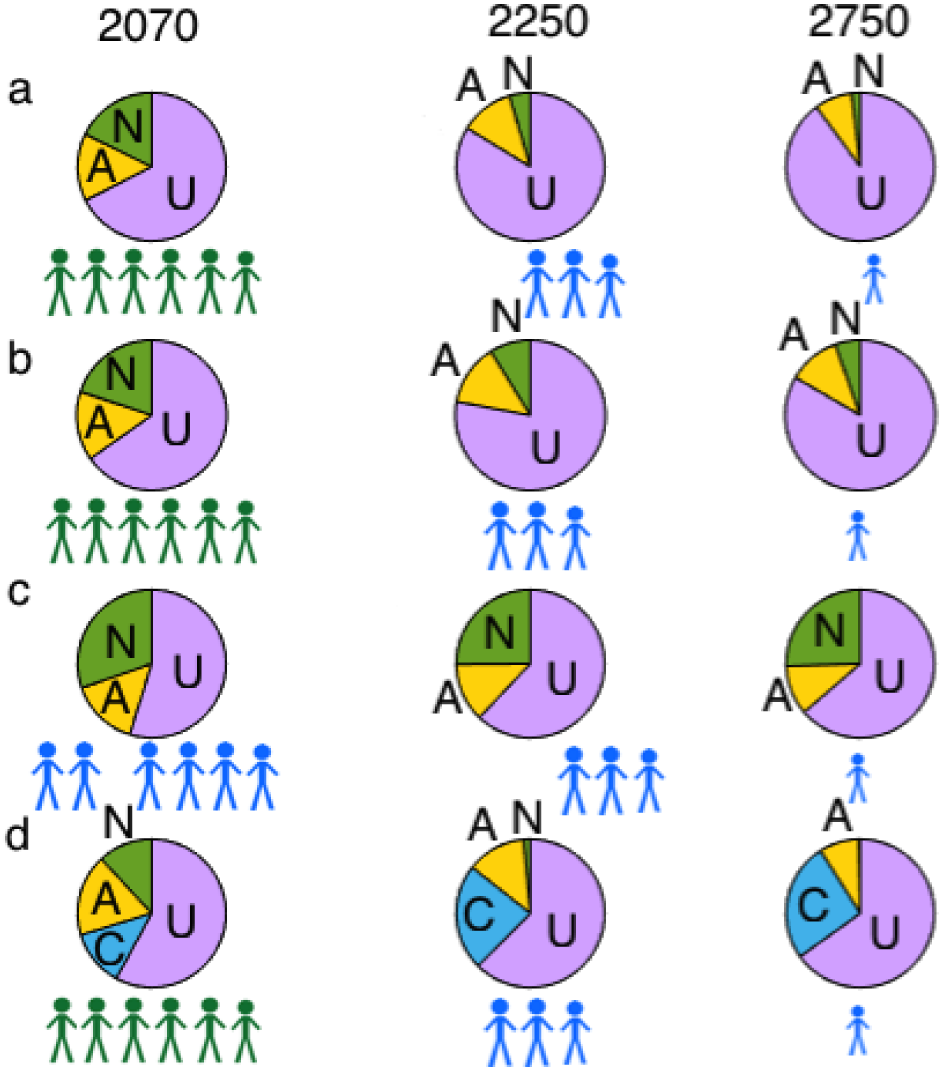
One-region (high-tech), all scenarios – land and population patterns. a) BAU – In the one-region scenario, there is no dispersal of goods or people, but there is still inequality. When technological development is rapid, all individuals have a good (green) or excessive (blue) well-being. The population size is smaller than two-regions (*P* = 7.5 in 2070, *P* = 2.8 in 2250, *P* = 0.4 in 2750). b) No status – There are no qualitative changes to the human population dynamics. There is however more natural land (*N*) and agricultural land (*A*) throughout the simulations. c) Restoration – Restoration increases human well-being, maintaining excessive well-being throughout the simulations. There is over 3 Bha of sustained *N* and 1.4 Bha of sustained *A*. The population size (*P*) is similar to the BAU scenario, with more resources, hence the greater well-being. d) Conservation - Conservation has similar impacts on land dynamics to restoration, yet *N* is converted into conserved land (*C*), which cannot be used by individuals. The population is low and therefore can be maintained by the services from *C* and *A*.

### 3.6 Status scenario

Lastly, in a scenario in which individuals are not allowed to change income status, thereby keeping access to resources limited in lower income populations (in both HI and LI regions), has little impact on the results. The number of individuals in each subpopulation changes; however the population size, by region, remains the same (Fig. A2). The difference in wellbeing is slight, yet there are no qualitative changes to the results. As there is no change in population dynamics, the land cover remains the same compared to the BAU scenario.

### 3.7 Comparison with the one-region system

The one-region system is much more stable than the two-region system. There are fewer feedbacks in the one-region system, which means the outcome of each action is more deliberate and achieves the desired goal. For example, the one-region case with restoration shows that restoration of natural land improves well-being (Fig. 5c), in addition to sustaining natural land at *N* = 3.3 Bha.

Unlike the two-region system, the one-region system maintains natural land (*N*), smaller populations, and a consistent state of well-being for all scenarios. The one-region system is strongly influenced by the rate of technological growth. Fast technological growth leads to a higher income scenario with excessive well-being, whereas, slow technological growth leads to mostly poor well-being populations with less than 3B individuals globally (Fig. A3).

Status change makes no qualitative difference (Fig. 5b). The population is all higher income already, so preventing the movement of individuals between income groups has little impact. The long-term population and land projections of our model are not necessarily realistic predictions, but they give an indication of the trends that can be expected for business as usual practices and alternative scenarios. Who is using what resources and in which regions has a major impact on the outcome of the business as usual model and the alternative scenarios. People and land-use shape recruitment, mortality and dispersal patterns.

## 4 CONCLUSIONS

The complex interactions between land, people and technology, make it difficult to predict the success of sustainable management policies. For example, restoration has the potential to promote higher sustained populations with improved well-being. However, we find that the multiple feedbacks between dispersal and resource accessibility are driven towards growth at the expense of well-being, as individuals move to where resources are more accessible and growth is possible. However, this eventually exhausts all the resources, leading to few accessible resources, a massive population, and a poor well-being.

In all model scenarios, it is evident that technology in higher income regions provides an advantage in the population’s ability to access resources and disperse, consequently contributing to the poor well-being of those less fortunate. Inequality in society has not always been as marked as today, early societies are generally recognized for their egalitarian social structures. Inequality within the society is thought to have emerged at the beginnings of agriculture (Price, 1995), or maybe as early as the Upper Paleolithic Era (Hayden, 2001). In recent years, inequality has been rising, which alters the access to resources and ultimately impacts the average global well-being. In our model, there are multiple layers of inequality in the global, two-region system – differences in technological development, education, infrastructure etc.– which reinforce power dynamics and keep the higher income population thriving, often at the expense of the lower income population. It was not possible in the scenarios we evaluated to have equal technological development in both regions. Inequality is a major impediment to sustainable development and improved well-being. From the model we can conclude that any effort to reduce land degradation, promote conservation or implement natural land restoration first needs to ensure adequate access to resources for all. There will always be inequality, but policy-makers should focus on reducing the gap, as inequality not only threatens the global societal well-being, but also impacts the environment and development, both locally and globally. We have just touched the surface of inequality here. Future work will take a more complete look at inequality and differences in consumption.

The one-region case describes a system where neither people, nor resources can disperse. This hypothetical system gives a glimpse into a world with reduced inequality and greater resource accessibility for all individuals. The one-region case maintains consistent well-being and results in slower depletion of agricultural and natural resources. Moreover, the land management scenarios simulated in the one-region environment indicates more or less the desired goal of each land action. Showing that the movement of people and goods can undermine well-intended actions, and can lead to confusion or dissociation with the land. In the one-region model simulations, there is still a degree of inequality, when it comes to individual access to resources, yet the technological and social development are the same, which removes many layers of inequality from distribution or dispersal, and increases sustainability.

That is not to say dispersal should be limited, as there are numerous benefits to human migration, such as technological development, economic stimulus and cultural diversity (Damelang and Haas, 2012). There are also numerous social factors to consider that are beyond the scope of this paper. Simply, the fact that individuals can move and make decisions based on resource accessibility, necessitates more forethought when it comes to land policies, and consumption practices. Dispersal plays a major role in undermining policies and conservation in our model, by masking feedbacks from the environment and delaying sustainable practices. Therefore, it is crucial to gain a better understanding of migration behaviours, the motivations for migration and how individuals adapt to their new environment.

The business as usual scenario provides a grim outlook on human well-being. Natural land conservation is one potential avenue for improving the long-term well-being of the human population and the natural environment; however, land patterns are strongly interlinked with social patterns. Dispersal in the model is driven by the amount of natural land and conserved land, as a proxy for greater ecosystem services and higher well-being. The extent of this influence may be over emphasized in the model. We are unable to conclude that these are realistic patterns of movement with conservation, but it does raise further questions about spatial interactions between people and nature. This suggests that the success of conservation in our current global system, with inequality, migration and trade of resources is highly susceptible to the spatial dynamics of society.

We are a global society with different land use patterns, social inequality, and the movement of people and goods. The spatial aspect of a global system with two distinct regions, allows for faster development of technology, higher peaks in population size, and generally speaking lower population well-being. The unequal access to resources and differences in technological progress, including the development of social structures, education and infrastructure, alter the outcome of natural and agricultural land sustainability and social policies. These scenarios do not include further degradation of natural land or agricultural land by way of climate change, changes in consumption, disease or civil unrest. We only look at the feedbacks between technology/innovation, human population dynamics and land cover. Even without such stochastic events or secondary effects, the scenarios show the rapid degradation of land and the counter-intuitive impact of well-intended policies. The potential for stochastic events to perturb the system could be enormous, considering the negative outcomes in a relatively ideal system. Future work will elaborate on the impact of land management and social equality on global socio-ecological sustainability.

## 5 ACKNOWLEDGMENTS

## Appendix

### Model in detail

The model represents a simplified global system with two regions classified by their income level (*j* = *L, H*) and each with two subpopulations (*i* = *L, H*). Individuals are categorized as either lower or higher income subpopulations, within higher or lower income regions (*P*_*i, j*_, *i* = *L, H, j* = *L, H*), similar to the model by Cumming and von Cramon-Taubadel (2018). The two regions are composed of natural land (*N*_*j*_, *j* = *L, H*), agricultural land (*A*_*j*_) and unused land (*U*_*j*_), which is the total land area (*L*_*j*_) minus *A*_*j*_ and *N*_*j*_. Land-use practices include local (*j*, such that *j* = *L, H*) and foreign (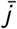, such that if *j* = *L*, 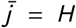 or *j* = *H* and 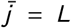) use of land, including functions describing degradation (*dN*_*j*_ (*P, T*), *dA*_*j*_ (*P, N, C*)), cross-degradation (*xdN*_*j*_ (*P, T*), *xdA*_*j*_ (*P, N, C*)), conversion (*cv*_*j*_ (*P, T*)), restoration (*r t* _*j*_ (*P, N, A, C*)) and options for conservation (*cs*_*j*_ (*N, C*)).

The human population is able to change income status within their own region given by the functions *s*_*i, j*_ (*P, N, A, C, T*) and 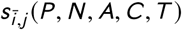, where *i* is the population in either the HI or LI subpopulations and 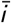 is the opposite of *i*. Individuals are able to disperse from one region to another, given by the functions *δ*_*i, j*_ (*P, N, A, C*), *T* and 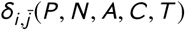. The growth of each subpopulation varies significantly with their access to resources, which is described by the function (*g*_*i, j*_ (*P, N, A, C, T*)).

The higher income region initially has a higher overall income level, such that those in the lower income subpopulation in the higher income region have a greater access to resources than individuals in the lower income region. Similarly, the technological development in the higher income region is initially faster than the lower income region. However, all the equations are dynamic and have the ability to change. The rates of change are given by the following differential equations:

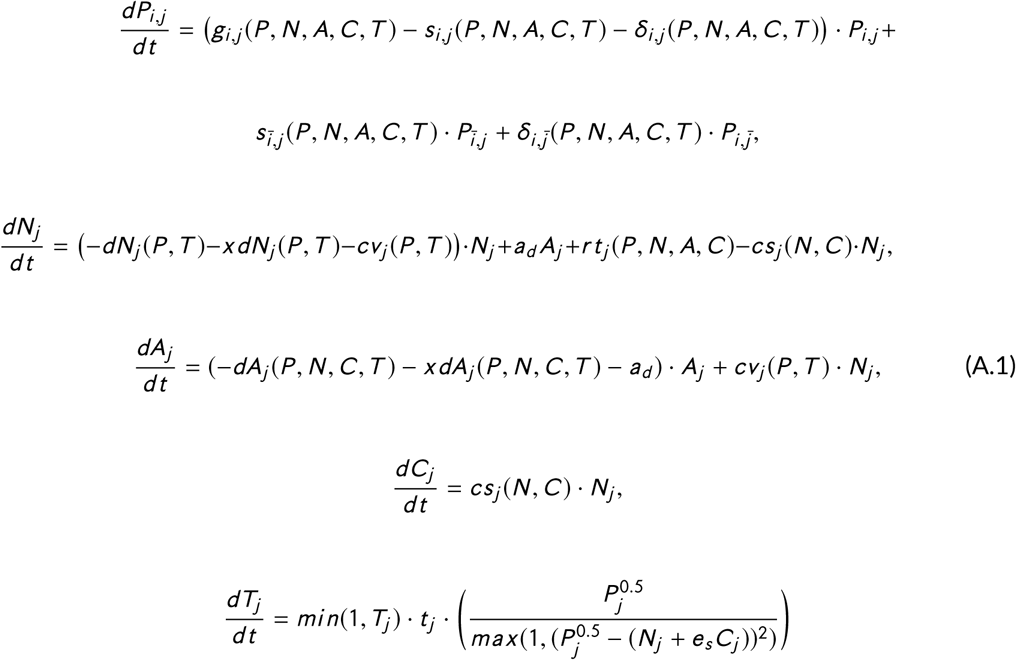

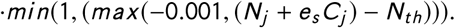

### Human population

Population growth rate (*g*_*i, j*_) is determined from the non-monotonic curve relating recruitment and mortality rates to resource accessibility (*R*_*i, j*_). The details of this theory are described in Henderson and Loreau (2019). The recruitment and mortality equations are modified to incorporate distinct regions, as such the resource accessibility units are given in ha/ind. rather than Bha, as is described in Henderson and Loreau (2019), which requires a calibration of the recruitment equation to maintain the characteristics of the curve. The curve shape is the same in both models, however the range is truncated here, which results in a sharper rise and decline for each hectare of accessible resource.

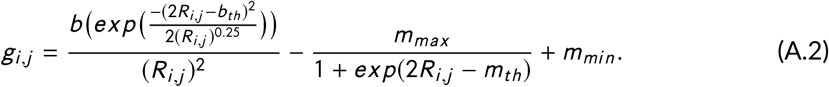

The maximum recruitment rate is given by *b*. There is a transition in social norms from survival mode (i.e., quantity) to thrive mode (i.e., quality) which occurs when resources reach a certain threshold of accessible resources (*b*_*th*_). The mortality rate is also threshold dependent (*m*_*th*_), such that there is a sharp drop from maximum mortality (*m*_*max*_) when *R*_*i, j*_ reaches *m*_*th*_. Unlike recruitment, mortality cannot equal zero, therefore *m*_*mi n*_ represents the minimum mortality rate.

Once a certain amount of accessible resources 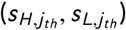 are obtained, an individual can change status from higher income to lower income (*s*_*H, H*_ (*P, N, A, C, T*), *s*_*H, L*_ (*P, N, A, C, T*)) or from lower to higher income (*s*_*L,L*_ (*P, N, A, C, T*), *s*_*L,H*_ (*P, N, A, C, T*)).

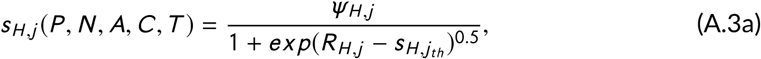

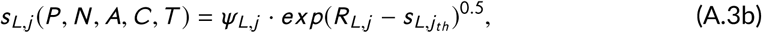

where *ψ*_*i, j*_ is the status shift coefficient that creates a bias in status change. Status changes with the resource accessibility (*R*_*i, j*_); however inequality in our social system makes it more difficult for lower income individuals break the poverty cycle (Payne, 2005). What we consider higher income occurs at *R* = 2ha/ind.; however, it is more difficult to become higher income, therefore 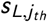 is set at 2.5. When access to resources drop below 1.5 in the model, individuals have a moderate well-being, which is considered to be the case for lower income individuals, therefore 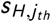 is set at 1.5; once the resource access for individuals in the HI subpopulation dips below 1.5 ha/ind. the individual becomes lower income. The values for 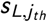 and 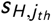 were estimated from the global ecological footprint average for each income group (details provided below in Population calculations and groupings section). Individuals are able to move from one region to another by comparing the resources accessible in the other region 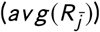 compared to their own resource accessibility (*R*_*i, j*_), based on their income status (*i, i* = *H, L*) and region (*j, j* = *H, L*). Individuals disperse to the region with greater resource accessibility given by the function:

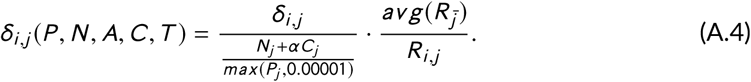

The equation for dispersal was modified from Keegan (1995), where movement is a logistic equation reflecting the relationship between births, deaths and resources; and Potapov et al. (2014). Here, a sigmoidal curve is used to represent the relationship between resource accessibility and dispersal.

Not all individuals are able to disperse at the same rate, inequality and policies limit dispersal, therefore the rate of dispersal (*δ*_*i, j*_, where *i* = *L, H*, subpopulation income status, and *j* = *L, H*, the region) is different for each subpopulation. Individuals also take into consideration the fraction of natural land (*N*_*j*_) and conserved land (*C*_*j*_) per person (*P*_*j*_). More natural and conserved lands presumably mean greater well-being. *α* changes the influence of conservation on migration.

### Technology

Technology presents a large unknown in terms of future potential. Generally, technology growth is represented by a sigmoidal curve in the literature (Henderson and Loreau, 2019). In the majority of cases, the technology output described here follows a sigmoidal trend (Fig. A1), but instead of imposing a threshold, it is the population density and natural land cover (both *N* and *C*) that determine the threshold, which allows more dynamic output and variations in onset, steepness and duration of the technology transition. In the framework outlined by Galor and Weil (2000), technological progress depends on population size and human capital. We argue that it is more plausible to assume that technological change depends additionally on population density, as population density facilitates communication and exchange, increases the size of markets and creates the required demand for innovation (Klasen and Nestmann, 2006).

We used historical data to calibrate the higher and lower income technology variables and differential equations. Technology growth rate up until present should lie between economic growth rate, which has grown at the same pace (Motesharrei et al., 2016) and technological growth, measured by advances in computer processing and inventions, which is 10 times faster than human population growth.

In the model, technology is a major driver of population dynamics via resource accessibility. It is estimated that higher income regions are more developed than lower income regions, in terms of education, medicine, machinery, etc. (The World Bank, 2019a). Therefore, we include two technology variables, one for each region (*T*_*j*_, *j* = *L, H*) with different growth rates (*t* _*j*_). The technology growth curve is a function of population size (i.e., *P*_*j*_ = *P*_*H, j*_ + *P*_*L,j*_, where *j* is the region, density (i.e., 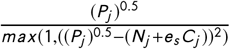), and natural resources (i.e., *N*_*j*_ and *C*_*j*_). Technology has been shown to increase with population density, however there becomes a point where the number of individuals exceed the capacity of natural land (*N*_*th*_) (Clow, 1998). Conserved land provides ecosystem services, minus provisioning services, therefore only a fraction is useful for technological development, given by *e*_*s*_.

Technology is a major determinant of power and resource accessibility – determining who will use what land, when, and how. We assume that technology builds upon itself, therefore the region with more technology has the potential to develop new technologies more quickly, hence the term *min*(1, *T*_*j*_).

Technological innovation does not create new human capabilities and production process, it only finds new ways to tap into and harness existing natural processes and energy flows (Clow, 1998). To continue economic growth indefinitely, technological innovation has to continue on a coordinated and indefinite basis. If not, economic expansion will place greater demands on the environment and cause more ecological disruption; as a result, one quickly runs into limitations on production arising from the inability of the Earth to supply resources and waste absorption. Therefore, we assume there is an ecological threshold in the development of technology (*N*_*th*_, eqns. A.1 & A.6). There is no such thing as indefinite improvement in technological efficiency or indefinite ability to tailor ecosystems to deliver more resources or absorb and recycle more wastes.

In terms of cost-benefit analysis, many have suggested that technology and resource prices can only prolong resource accessibility within certain environmental and social limits (Mudd, 2010; Prior et al., 2012; Schandl and West, 2010). Hence, the use of a density-dependent technology curve (fig. A1). Below is a simplification of the density- and nature-dependent technology equation.

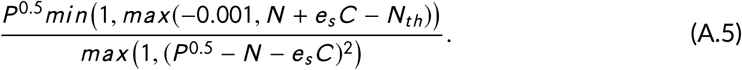

### Resource acquisition

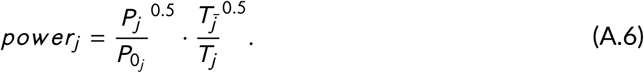

Power (*power R*_*j*_) refers to the share of resources the population, in each region, (*P*_*j*_) can access. Foreign access to resources is simply 1 − *power R*_*j*_. For example, higher income regions exert more power, therefore control the majority of land in their own region (*j*), yet still have access to a significant share in the other region 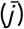. This share of land is given by the power ratio equation:

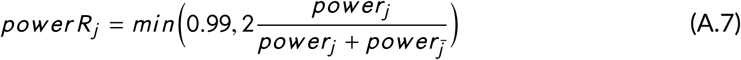

**FIGURE A1.**
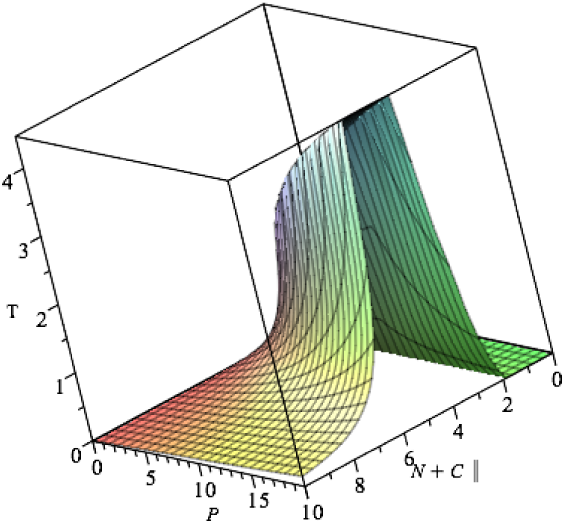
Non-monotonic technology curve that is density-dependent 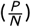 and nature-dependent (*N, C*).

The power dynamics for agriculture are the same as for natural land.

Resource accessibility per person is a function of the resources available (*N*_*j*_, *A*_*j*_) and how much access each individuals has to these resources through power and technology. It is assumed that technology (*T*_*j*_) determines the yield and ability to acquire the available resources.

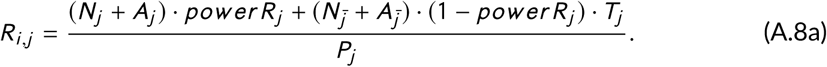

The power dynamics can drop the resource accessibility to unrealistic proportions. Therefore, when *R*_*i, j*_ < 1 the resource accessibility is scaled according to the famine state (*R* = 0.55), resources (*N*_*j*_, *C*_*j*_, *A*_*j*_), technology (*T*_*j*_) and people (*P*_*j*_).

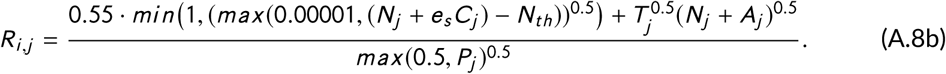

For the minority subpopulation in each region (when *i* ≠ *j*), we assume an unequal access to resources. Despite the same technology and development, the access is not distributed equally throughout the region. We scale the resource accessibility as follows, *R*_*L,H*_ = *R*_*H, H*_ · 0.65 and *R*_*H, L*_ = *R*_*L,L*_ · 1.6, according to GINI data from OECD (2019); The World Bank (2019b).

### Land dynamics

The composition is classified given FAO data on the eleven global land cover layers. Agricultural land (*A*_*j*_) is composed entirely of croplands, which occupies 12.6% of current land cover. Natural land (*N*_*j*_) describes the broadest range of land cover, including grasslands, herbaceous vegetation, mangroves, shrub-covered areas, sparse vegetation, tree-covered areas, for a total of 59.3%. Finally, the unused land describes the remaining land cover, artificial surfaces, bare soils, snow and glaciers, and inland water bodies (28.1%) (Food and Agriculture Organization of the United Nations, 2019).

### Natural land

The degradation and consumption of natural land in each region (*j* = *L, H*) depends on the population size within each region (*P*_*j*_), the demand and degradation of resources (*dN*_*j*_) and the proportion land under control of population (*power R*_*j*_), *P*_*j*_.

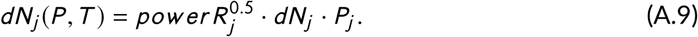

In both, degradation and cross-degradation, the square root of power is used to determine the proportion of land used by each region, as the impact of degradation can extend beyond the land under question and percolate to other patches (Kun et al., 2019). Furthermore, the ecosystem services from one area are beneficial to the adjacent areas, therefore reducing the natural land area can reduce the resilience of the surrounding area. For example, pest control in wheat crops benefits from natural predators and as Woodcock et al. (2016) explain the spill-over of natural pest control services declined with distance from the crop edge.

The remaining proportion of land not used by the local individuals (1 − *power R*_*j*_) gets consumed and degraded by the individuals from the other region (*P*_*j*_). In what is termed cross-degradation:

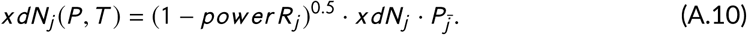

The coefficient for cross-degradation and consumption, *xdN*_*j*_, is assumed to be smaller than *dN*_*j*_. The values assigned to the coefficients are given in Table A1.

Conversion of natural land (*N*_*j*_) to agricultural land (*A*_*j*_ is necessary to supply the current and future populations with adequate food. The equation is given by

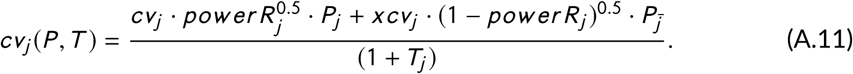

Agriculture is part of the global trade network and as such agricultural production in one region is likely to be consumed by another region. Conversion depends on the demand from each region *cv*_*j*_ (local) and *xcv*_*j*_ (foreign) and what share of the land each region is able to manipulate (*power R*_*j*_ and 1 − *power R*_*j*_). Furthermore, technology allows the yield to improve without increasing land cover. Hence, when technological development (*T*_*j*_) increases, conversion (*cv*_*j*_ (*P, T*)) of *N*_*j*_ to *A*_*j*_ declines.

Restoration is both the natural and human process of converting unused or degraded land (*U*_*j*_) back into natural land (*N*_*j*_).

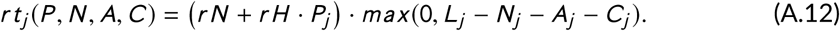

The rate of natural land restoration (*r N*) is the inverse time required to return *U*_*j*_ back into a natural space. Human land restoration is another model scenario, used to analyse the active process of returning degraded land to a natural state. Therefore, *r H* is the constant rate of restoration per person (*P*_*j*_). A minimal restoration is expected in the business as usual scenario, such that *r H* = *r N*.

Conservation in the model refers to a fraction of land set aside for non-provisioning ecosystem services.

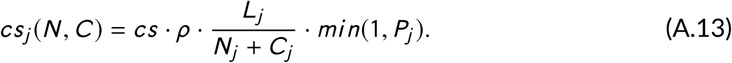

The goal of conservation is to maintain a certain proportion of natural land (*C*_*j*_). Therefore, natural land (*N*_*j*_) is set aside and continually maintained, so long as there are people to govern the conserved land, *min*(1, *P*_*j*_). The conservation of natural land (*C*_*j*_) is a model scenario, where a proportion of land (*ρ*) is set aside and maintained at a rate of *cs* · *L*_*j*_ /(*N*_*j*_ + *C*_*j*_).

### Agricultural land

The majority of newly converted agricultural land (*A*_*j*_) area is derived from natural land (*N*_*j*_), leading to degradation of *N*_*j*_. It has also been shown that agricultural production improves with surrounding natural land area and that degradation is intensified when there is a lack of supporting ecosystem services (Bennett et al., 2009; Braat and De Groot, 2012). Therefore, agricultural degradation is a function of the degradation rate (*dA*_*j*_) and ecosystem service influence:

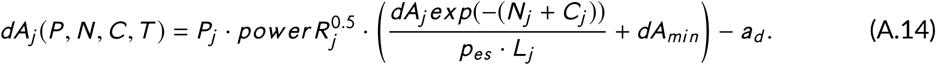

Degradation is impacted by regulating services, it has been shown that natural land provides essential assets that increase the yield, diminish pests and provide numerous benefits to agricultural land (Swinton et al., 2007). Therefore, degradation is intensified when the natural land (*N*_*j*_) and conserved land (*C*_*j*_) area do not represent the proportion *p*_*es*_ of the entire land area. From studies on the benefits of pollination (Morandin and Winston, 2006) on agriculture and improved insect diversity and pest control in complex agriculture systems (Bianchi et al., 2006; Söderström et al., 2001), the required proportion of land for the flow of ecosystem services and production (*p*_*es*_) is estimated at 0.3 of the region (*L*_*j*_). As the population (*P*_*j*_) grows, so does the consumption of agriculture goods and degradation. The regional population exerts a proportion of the degradation, which is represented by *power* 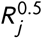. Even if there are sufficient supporting services there will still be a minimal rate of agricultural degradation 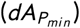, as a result of consumption and modern agricultural practices and urbanization (Azadi et al., 2011; Southgate et al., 1990; Smetanová et al., 2019). In addition to human degradation of agricultural land, natural degradation occurs as a result of soil erosion at a rate approximately 3 to 8 times slower than human-caused soil erosion, hence *a*_*d*_ is set to 0.0001/yr (Nearing et al., 2017).

Agricultural land is further consumed and degraded by the foreign population 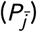 in the region *j* (*j* = *L, H* ; 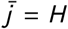 if *j* = *L* and 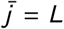 if *j* = *H*). Cross-degradation is determined by the proportion of land used by foreign entities ((1 - *power R*_*j*_)^0.5^).

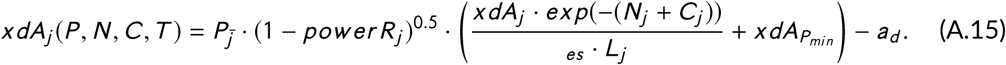

Cross-degradation shares the same characteristics as degradation, except the rate of degradation (*xdA*_*j*_) reflects the demands from the foreign region. The minimal cross-degradation (*xdA*_*Pmin*_) is also dependent on the demands from the foreign region.

**TABLE A1.** Description of model variables and parameters, including parameter values, initial conditions and units.

| Variables & Parameters | Definition | Value BAU | IC/Value Range | Units |
| --- | --- | --- | --- | --- |
| $P_{i,j}$ | Population subgroups ($i = L, H; j = L, H$) | — | $P_{0_{L,L}} =, P_{0_{H,H}} =, P_{0_{H,L}} =, P_{0_{L,H}} =$ | B inds. |
| $N_j$ | Natural land cover in each region ($j = L, H$) | — | $N_{0_L} = 6.4, N_{0_H} = 6.5$ | Bha |
| $A_j$ | Agricultural land cover in each region ($j = L, H$) | — | $A_L =, A_H =$ | Bha |
| $C_j$ | Conservation land cover in each region ($j = L, H$) | — | $C_{0_L} = 0, C_{0_H} = 0$ | Bha |
| $T_j$ | Technology in each region ($j = L, H$) | — | $T_{0_L} = 0.035, T_{0_H} = 0.085$ | NU |
| $R_{i,j}$ | Resource accessibility per capita in each subpopulation ($i = L, H; j = L, H$) | function | function | ha/ind. |
| $N_{th}$ | $N$ threshold for resource accessibility and technological development | 1.9 | — | Bha |
| $e_s$ | Fraction of conserved land accessible to inds. | 0.3 | — | NU |
| $b$ | Maximum recruitment rate | 0.058 | — | /yr |
| $b_{th}$ | Recruitment threshold causing demographic transition | 3.12 | — | ha/ind. |
| $m$ | Mortality rate | 0.03 | — | /yr |
| $m_{th}$ | Mortality rate threshold causing demographic transition | 2 | — | ha/ind. |
| $m_{min}$ | Minimum mortality rate | 0.004 | — | /yr |
| $s_{i,j,th}$ | Resources needed to change status for each income level ($i = L, H$) | $s_{L,j,th} = 2.5, s_{H,j,th} = 1.5$ | — | ha/ind. |
| $\psi_{i,j}$ | Status shift coefficient in each subpopulation ($i = L, H; j = L, H$) | $\psi_{L,L} = 0.005, \psi_{H,L} = 0.015,$<br>$\psi_{L,H} = 0.025, \psi_{H,H} = 0.025$ | — | /yr |
| $\delta_{i,j}$ | Dispersal rate for each subpopulation ($i = L, H; j = L, H$) | $\delta_{L,L} = 0.0005, \delta_{L,H} = 0.0008,$<br>$\delta_{H,L} = 0.0008, \delta_{H,H} = 0.001$ | 0 – 0.002 | /yr |
| $\alpha$ | Conservation factor on migration | 1 | 0.1–1 | NU |
| $dN_j$ | Local consumption & degradation of $N$ in region $j$ by subpopulations in region $j$ | $dN_L = 0.0005, dN_H = 0.0025$ | — | /ind · yr |
| $xdN_j$ | Foreign consumption & degradation of $N$ in region $j$ by subpopulations in region $\bar{j}$ | $xdN_L = 0.0025, xdN_H = 0.001$ | — | /ind · yr |
| $cv_j$ | Conversion rate from $N$ to $A$ in region $j$ by subpopulations in region $j$ | $cv_L = 0.0003, cv_H = 0.00095$ | — | /ind · yr |
| $xcv_j$ | Conversion rate from $N$ to $A$ in region $j$ by subpopulations in region $\bar{j}$ | $xcv_L = 0.0002, xcv_H = 0.00025$ | — | /ind · yr |
| $cs_j$ | Conservation rate from $N$ to $C$ in region $j$ | 0 | 0–0.001 | /ind · yr |
| $\rho$ | Proportion of $N$ to be conserved | 0 | 0–0.3 | NU |
| $rH$ | Anthropogenic restoration | 0.00001 | 0.00001–0.005 | /ind · yr |
| $rN$ | Natural restoration | 0.00001 | — | /yr |
| $p_{es}$ | Ratio of $N$ to $L$ that yields beneficial ecosystem services to $A$ | 0.3 | — | NU |
| $dA_j$ | Nature-dependent $A$ degradation rate in local region $j$ by subpopulations in region $j$ | $dA_L = 0.00075, dA_H = 0.0005$ | — | /ind · yr |
| $dA_{P_{min}}$ | Minimum consumption & degradation rate of $A$ in the absence of ecosystem services in region $j$ by subpopulations in region $j$ | 0.00005 | — | /ind · yr |
| $a_d$ | Natural degradation rate of $A$ | 0.0001 | — | /yr |
| $xdA_j$ | Foreign consumption & degradation rate of $A$ in the absence of ecosystem services in region $j$ by subpopulations in region $\bar{j}$ | $xdA_L = 0.0009, xdA_H = 0.0001$ | — | /ind · yr |
| $xdA_{P_{min}}$ | Foreign nature-dependent $A$ degradation rate in region $j$ by subpopulations in region $\bar{j}$ | 0.00005 | — | /ind · yr |

—, no other values were simulated; NU, no units.

**FIGURE A2.**
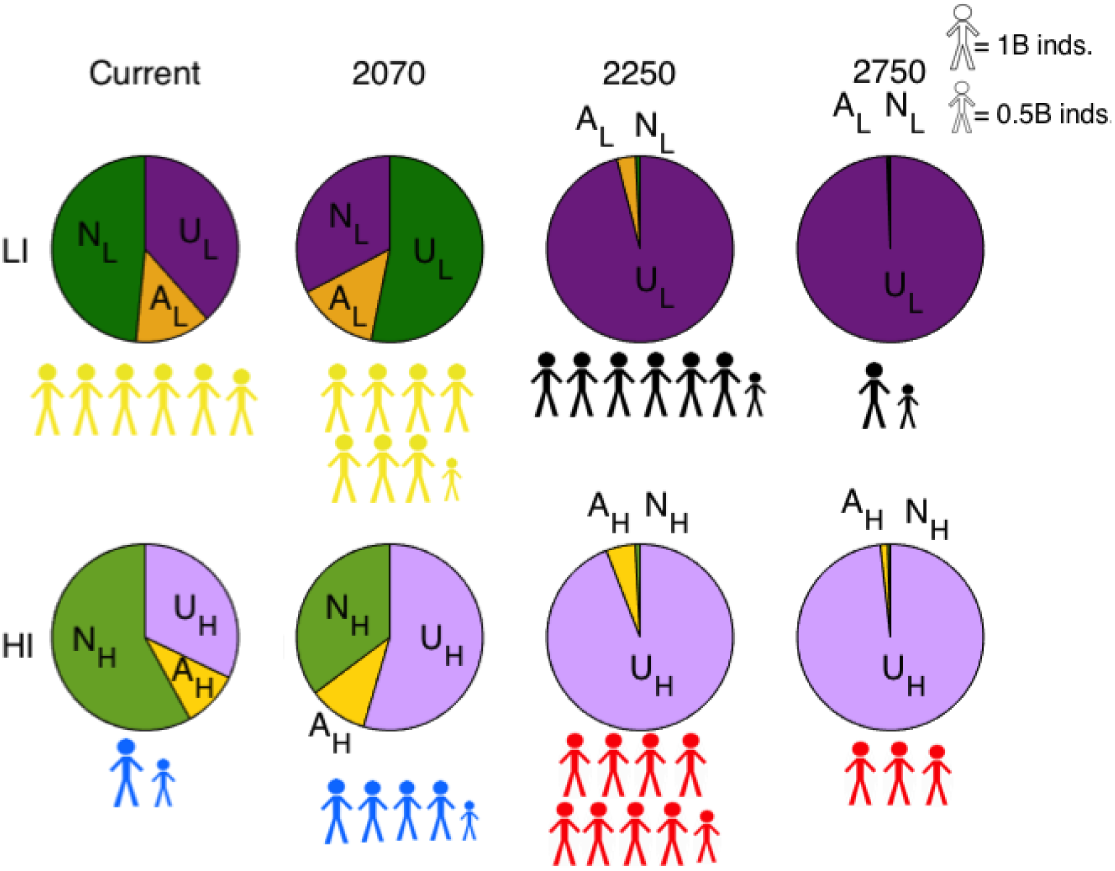
No status shift – land and population patterns. The land and population dynamics are unchanged from the BAU scenario. The population size (*P*_*L*_ and *P*_*H*_) are the same, the subpopulations are different, however that is not evident here. The well-being is qualitatively unchanged. The *P*_*L*_ transitions from moderate (yellow) to famine (black), while the *P*_*H*_ transitions from excessive (blue) to poor (red). The land cover is similar to the BAU scenario, where natural land (*N*_*L*_, *N*_*H*_) and agricultural land (*A*_*L*_, *A*_*H*_) are negligible.

### Additional results

#### Status

Globally, the population is becoming wealthier. Therefore, accelerating the movement of individuals through income status, allows more individuals to move to the higher income population, which decreases the number of individuals in the lower income population. This is often seen in rural-urban transitions, such that when individuals move from rural to urban regions, fertility decreases (Jensen and Ahlburg, 2004). However, in this scenario there is no change to the long-term resource accessibility or well-being to support the shift in population income levels, therefore the trend is similar to business as usual (BAU) scenario. In the case of no status shift (Fig. A2), the overall population in each region remains the same as the BAU scenario. In both cases, the collapse of resource accessibility is too strong to be overcome by shifts in income status.

#### Technology

The model suggests that technology is more than three times greater in the HI region than in the LI region. This is in range of the Human Development Index (HDI), where the countries with the greatest HDI (i.e., Norway, Switzerland, Australia) have HDI values that are 2.8 times greater than countries with the lowest values (i.e, Niger, Central African Republic) (United Nations Development Programme, 2019). The HDI considers education, income and life expectancy, all of which are considered components of our technology variable. Technology and population are intricately linked, which is evidenced through the multiple scenarios evaluated. Technological development is promoted by the number of individuals; however, when there are too many people and natural land is limited, this impedes technology growth. The non-monotonic resource curve (Fig. A1) suggests that technology can either help the population grow, if the well-being is in the moderate state or worse, otherwise the population decreases, if the population well-being is good or above, based on theory on the demographic transition (Henderson and Loreau, 2019).

#### Conservation

Conservation in either region, always benefits the HI population more than the LI population; however, conservation in the LI region encourages the development of technology and improves well-being compared to the business as usual scenario. Individuals in the HI region benefit from little competition for resources in their own region and take resources from the LI region, where the LI increase in technology staves off famine in the LI region.

#### Restoration

Restoration in a specific region (i.e., 1 of the 2 regions) results in similar trends to conservation in a specific region. Natural land and agricultural land are sustained in the region with restoration and the population disperses to the region with natural land restoration. In the higher income region, the population is very high (37B inds.) and the land area is minimal (*N* = 1 Bha and *A* = 0.2 Bha), causing a poor well-being in the region with restoration. In the lower income region, restoration causes unrealistically high population numbers (>200B inds.), as result of the positive feedback between people and technology, with adequate natural land. In both cases, the region without restoration becomes non-existent in terms of land and people in the long-term.

#### One-region

If the one-region system were to follow a low-tech scenario, what could be considered a world without the industrial revolution, the population would remain low and in a poor state (Fig. A3). Here land management policies can either help, minimally, or harm the population. Restoration provides slightly more resources to the population, which increases resource accessibility enough to reproduce more, but not enough to improve well-being (Fig. A3b). Conservation, initially decreases the well-being of the human population, by limiting the access to resources until the population declines sufficiently to reach a poor human well-being and sustainable land cover (Fig. A3d). In all the scenarios, some form of natural land is maintained, either conserved or accessible, and agricultural land area is consistent over the years.

**FIGURE A3.**
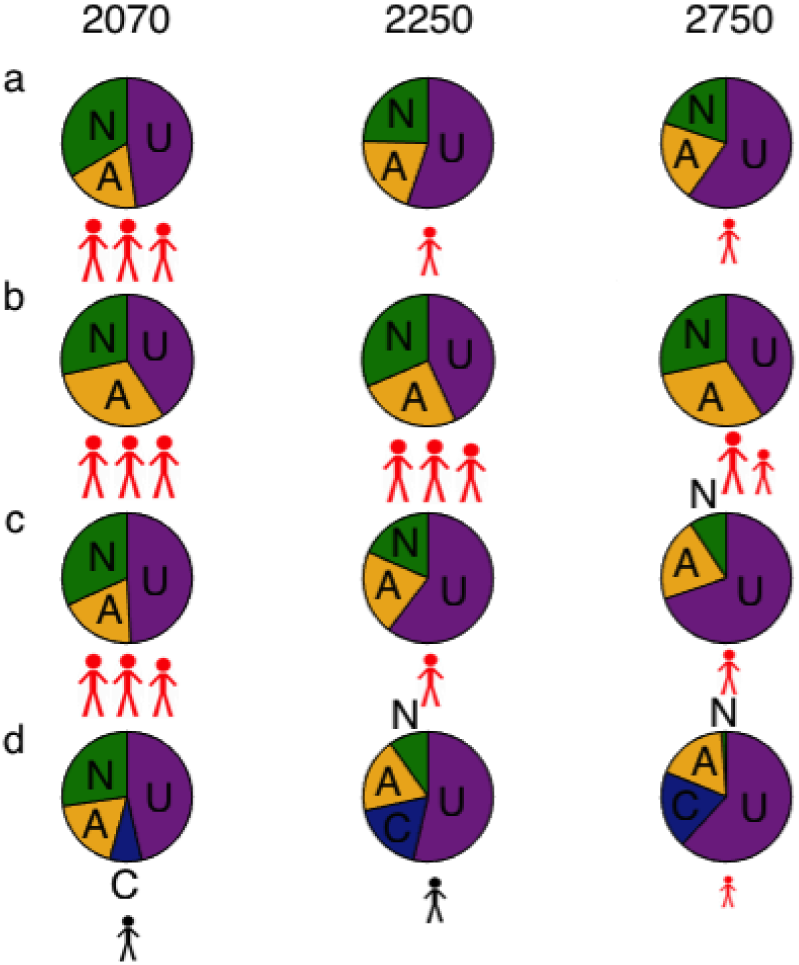
One-region (low-tech), all scenarios — land and population patterns. a) BAU - In the one-region, low-tech case, the population does not grow quickly, in 320 years the *P* increases by 1.8B. The slow growth is due to a lack of accessible resources, all individuals have a poor (red) well-being, except in the conservation scenario. The population size is between 4 times and 34 times smaller smaller than BAU two-regions (*P* = 2.8 in 2070, *P* = 0.5 in 2250, *P* = 0.3 in 2750). Significant fractions of natural land (*N*) and agricultural land (*A*) are maintained over time, yet *N* still declines (*N* = 4.3 Bha in 2070, *N* = 2.6 Bha in 2750) and *A* increases by 0.2 Bha. b) Restoration — A higher *P* is maintained over time, with restoration. There is also more *N* and *A* throughout the simulations. c) No status - The population size (*P*) and well-being (*W*) is similar to the BAU scenario. There is a decline in *N* from 4.1 Bha to 1.2 Bha, while *A* remains constant. d) Conservation - Conservation reduces the the population size (*P* < 0.3B inds.) and well-being (*W* = *f amine*), immediately after application. Once population declines to ≈ 0.1*B*, resource accessibility per person returns to *W* = *poor*.

### Data Analysis

#### Kernel Density Plots

**FIGURE A4.**
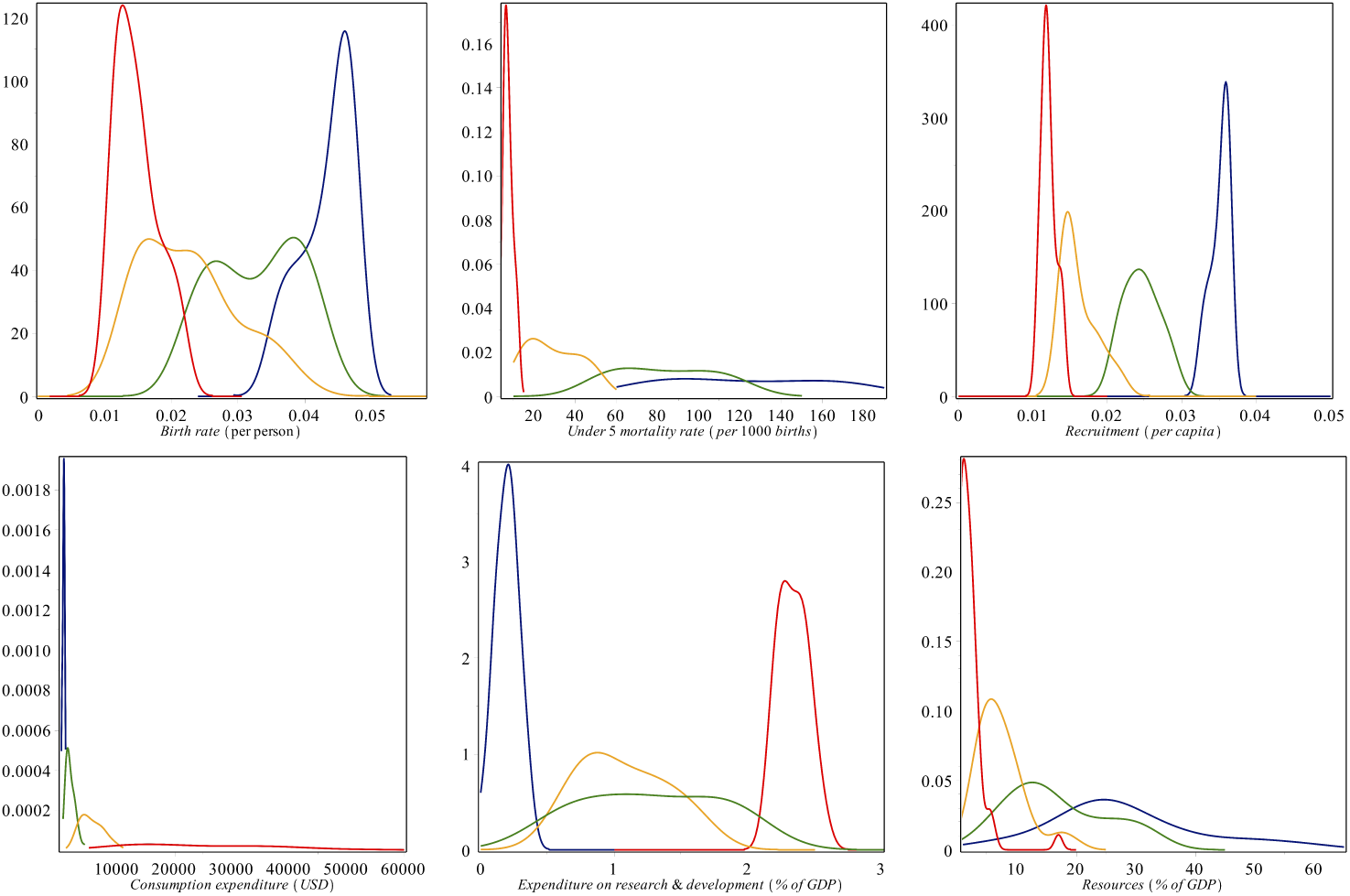
Kernel density plots to show the differences between high and low-income groups. The x-axis represents a global development indicator and the y-axis represents the calculated density. Each plot shows the range of values for each income group, as classified by the World Bank Group, over the last 60 years. There are clear clustering patterns that distinguish low-income countries from high-income countries, while the upper and lower middle-income countries lie in the middle and overlap with all income groups. The data for research expenditure in low-income and lower middle-income were sparse; therefore the lower middle-income group was generated using data for East Asia and Pacific (excluding high-income countries) and the low-income countries was calculated as an average of low-income countries with consistent data (Madagascar, Burkina Faso and Uganda). The plots were generated using data from the World Bank Group in Maple.

#### Population calculations and groupings

The information here was used to set the subpopulations. Data is from 2010 for population size and GDP for each country given by the World Bank Group (The World Bank, 2019a). Countries with a GDP per capita greater than 20 000 USD are classified as *P*_*H H*_ in our model; between 10 000 and 20 000 USD are classified as *P*_*LH*_ ; between 5 000 and 10 000 USD are classified as *P*_*H L*_ ; and below 5 000 USD are classified as *P*_*LL*_. These groupings were used as guidelines for the model simulations, for example the model simulations for population numbers were compared to the 2010 values to validate the model. These groupings also provided the basis for the model assumptions and kernel density plots. When compared to the World Bank Group classification, the bounds set in our model classification are higher on average than the World Bank, but the population within the regions of higher income and lower income remain the same.

We also include the ecological footprint of each country and an average footprint per income group. The ecological footprint is measured in global hectares, which includes water bodies. In our model we do not include water, therefore we reduce the resource accessibility (≈ ecological footprint) to account for the absence of water bodies. However, we maintain that *R*_*H H*_ ≈ 1.35 *R*_*LH*_ and *R*_*H L*_ ≈ 1.62 *R*_*LL*_, as indicated by the Global Footprint Network (2019).

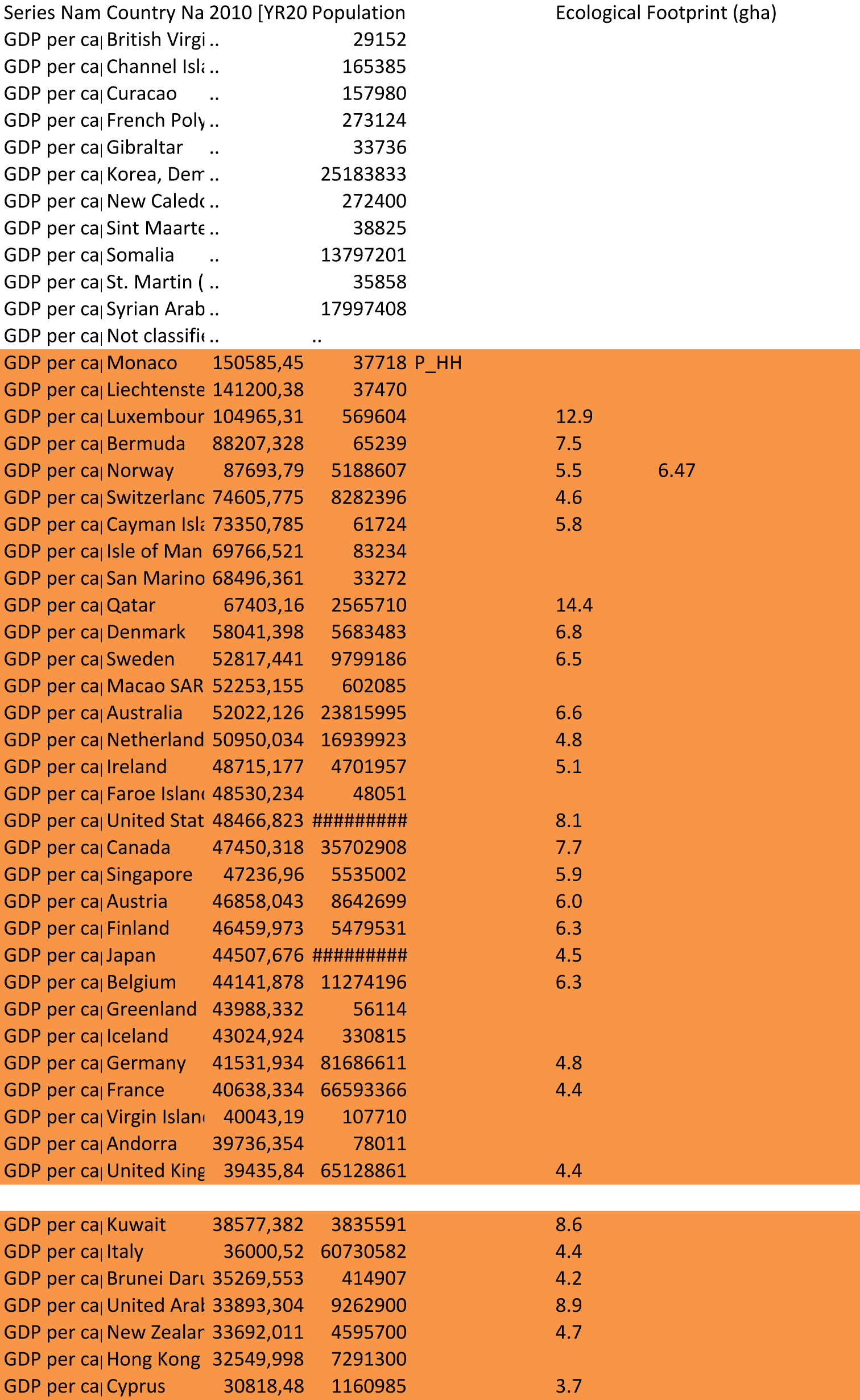
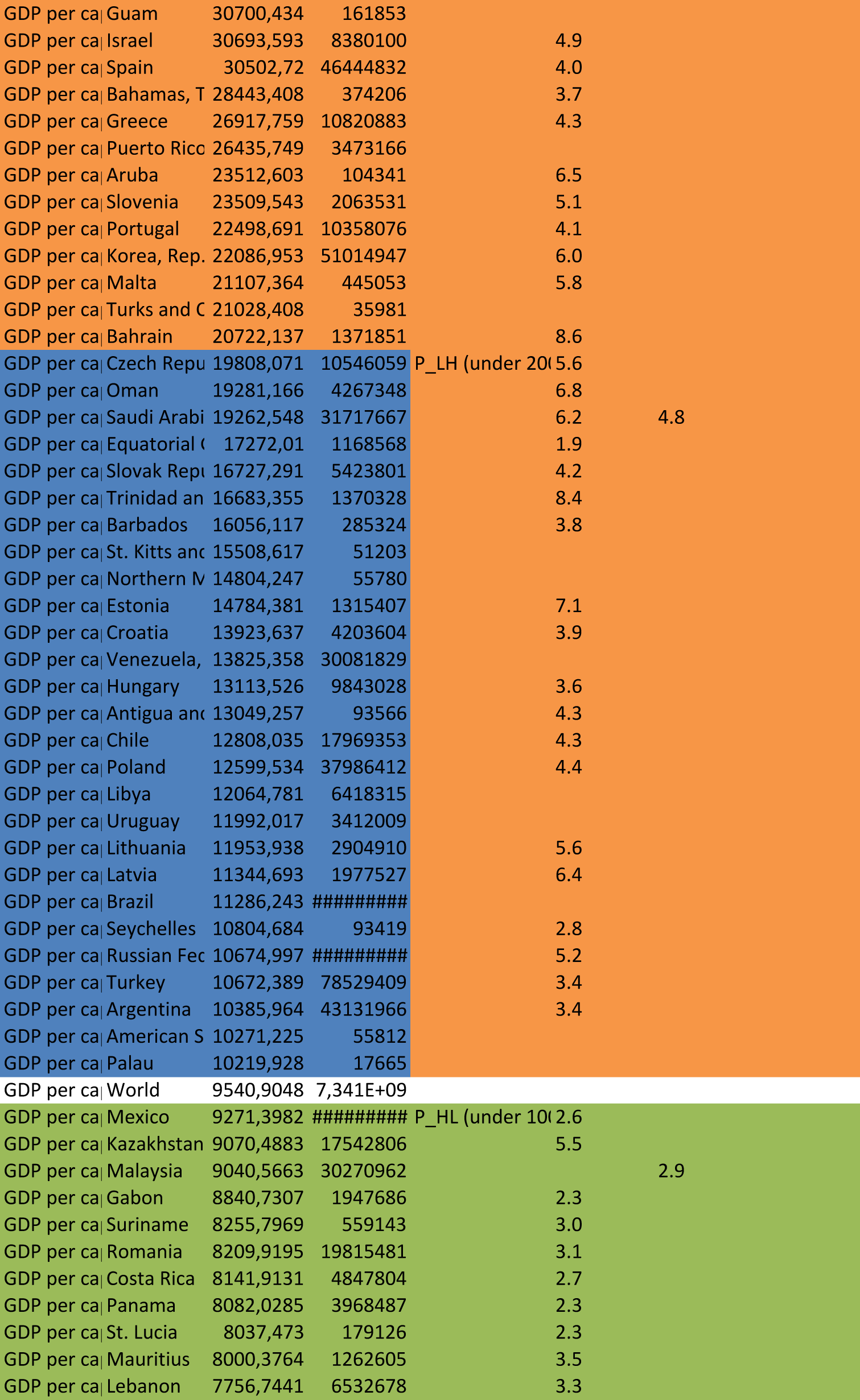
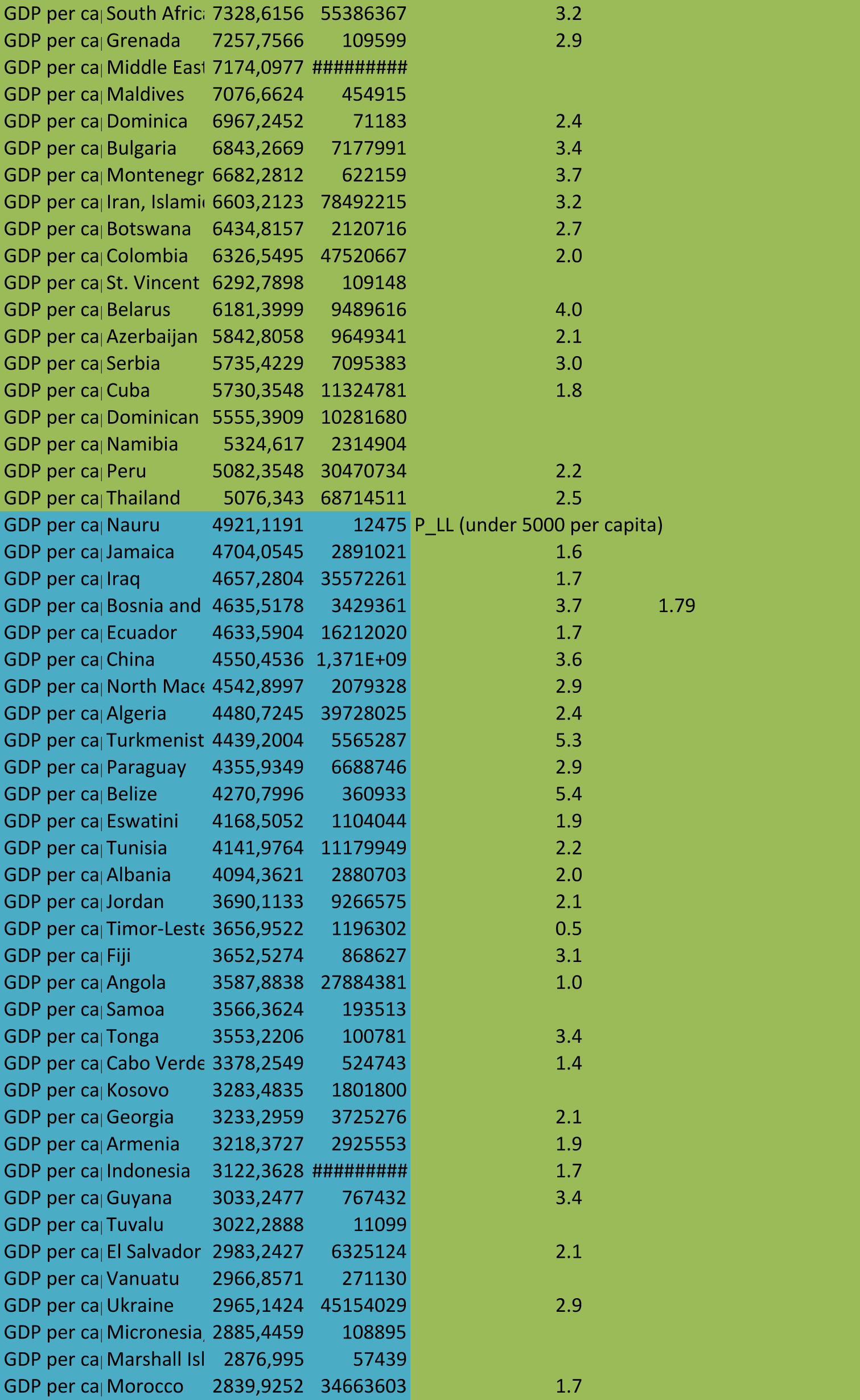
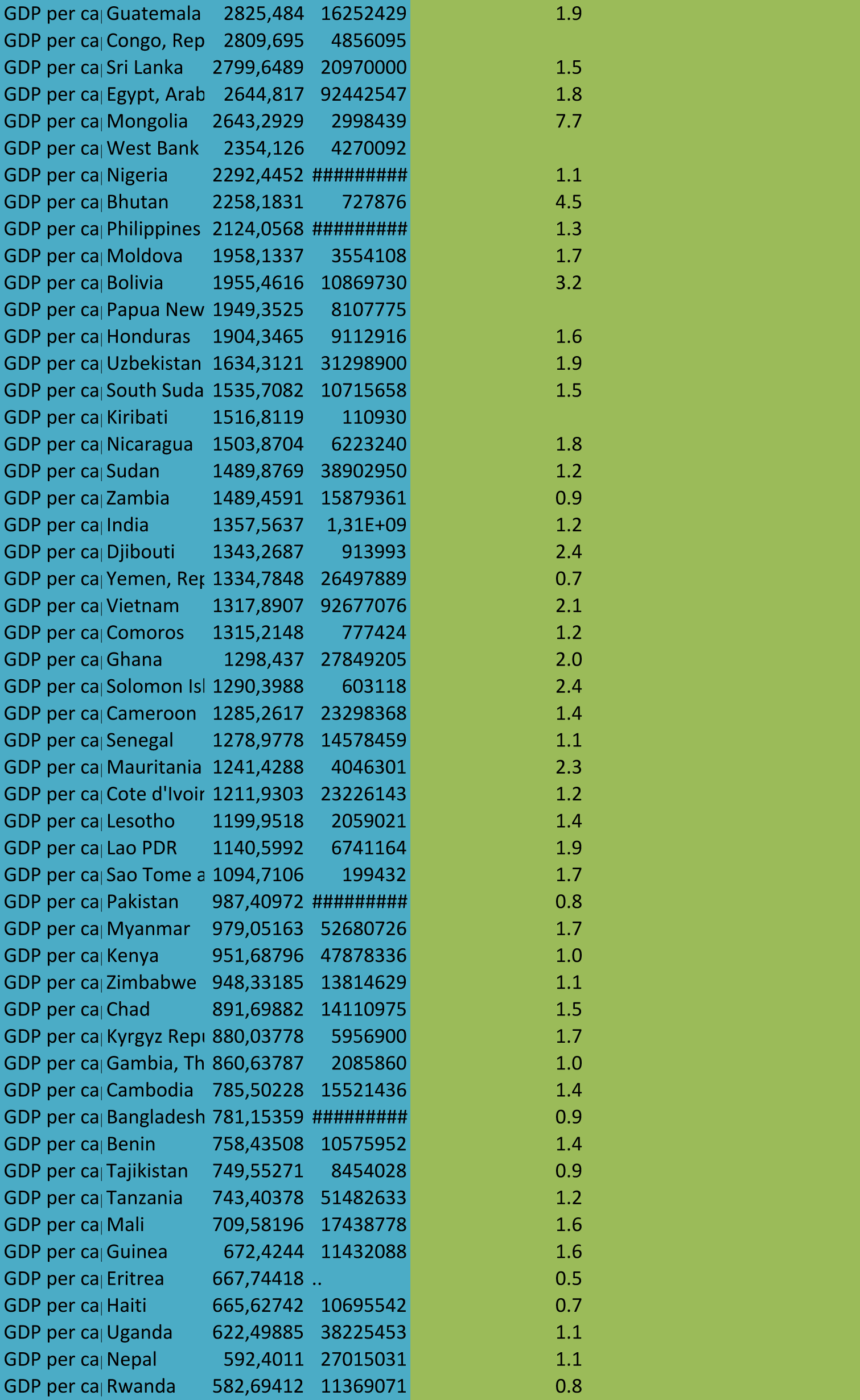
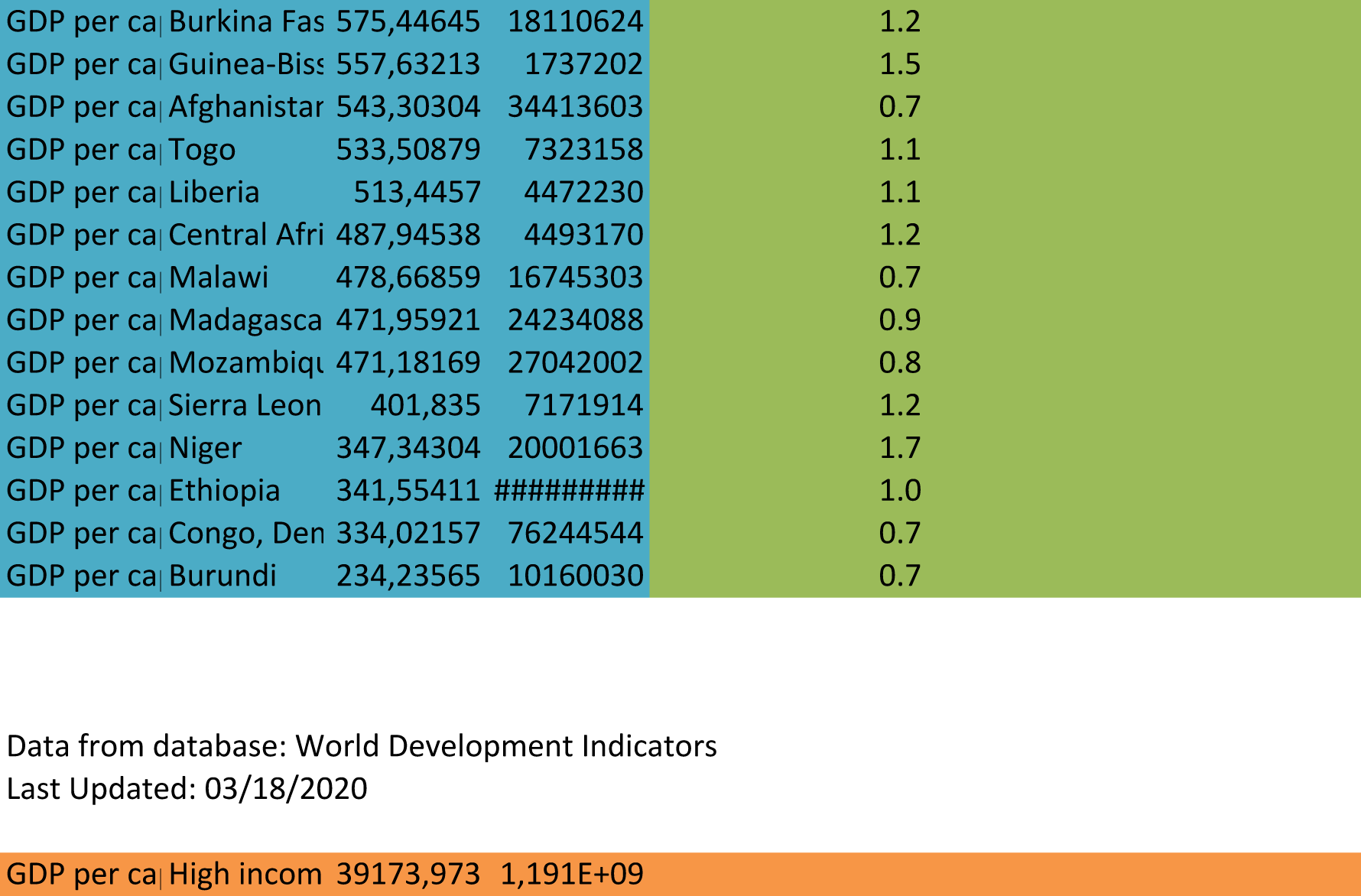

